# The Essential Role of O-GlcNAcylation in Hepatic Differentiation

**DOI:** 10.1101/2023.02.16.528884

**Authors:** Dakota R. Robarts, Manasi Kotulkar, Diego Paine-Cabrera, Kaitlyn K. Venneman, John A. Hanover, Natasha E. Zachara, Chad Slawson, Udayan Apte

## Abstract

**Background & Aims:** O-GlcNAcylation is a post-translational modification catalyzed by the enzyme O-GlcNAc transferase (OGT), which transfers a single N-acetylglucosamine sugar from UDP-GlcNAc to the protein on serine and threonine residues on proteins. Another enzyme, O-GlcNAcase (OGA), removes this modification. O-GlcNAcylation plays an important role in pathophysiology. Here, we report that O-GlcNAcylation is essential for hepatocyte differentiation, and chronic loss results in fibrosis and hepatocellular carcinoma.

**Methods:** Single-cell RNA-sequencing was used to investigate hepatocyte differentiation in hepatocyte-specific OGT-KO mice with increased hepatic O-GlcNAcylation and in OGA-KO mice with decreased O-GlcNAcylation in hepatocytes. HCC patient samples and the DEN-induced hepatocellular carcinoma (HCC) model were used to investigate the effect of modulation of O-GlcNAcylation on the development of liver cancer.

**Results:** Loss of hepatic O-GlcNAcylation resulted in disruption of liver zonation. Periportal hepatocytes were the most affected by loss of differentiation characterized by dysregulation of glycogen storage and glucose production. OGT-KO mice exacerbated DEN-induced HCC development with increased inflammation, fibrosis, and YAP signaling. Consistently, OGA-KO mice with increased hepatic O-GlcNAcylation inhibited DEN-induced HCC. A progressive loss of O-GlcNAcylation was observed in HCC patients.

**Conclusions:** Our study shows that O-GlcNAcylation is a critical regulator of hepatic differentiation, and loss of O-GlcNAcylation promotes hepatocarcinogenesis. These data highlight increasing O-GlcNAcylation as a potential therapy in chronic liver diseases, including HCC.

**Lay Summary:** Proteins in cells are modified by the addition of a single glucosamine sugar molecule called O-GlcNAcylation. Loss of O-GlcNAcylation in hepatocytes, the most common type of cells in the liver, causes the liver to lose its function and can result in increased liver diseases such as fibrosis and cancer.

**Graphical Abstract:** 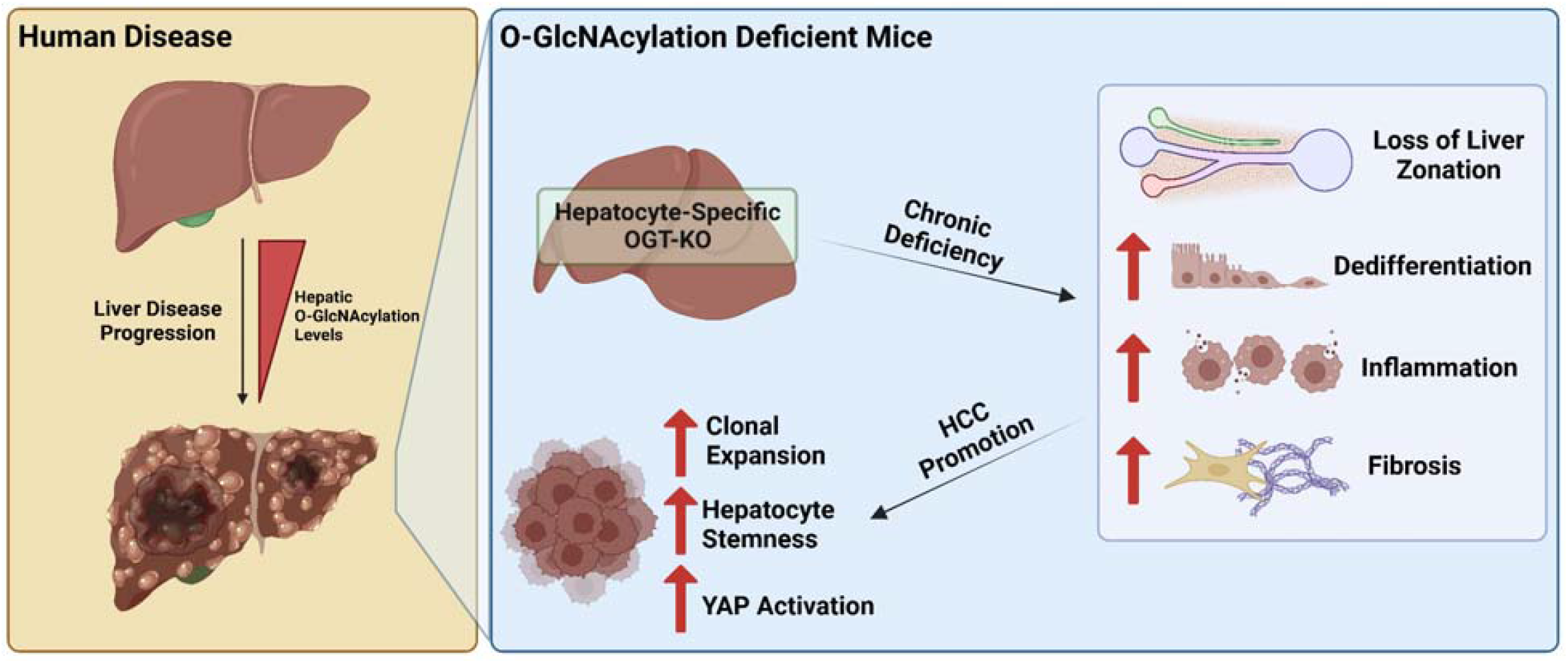

**Highlights:**

- Single-Cell RNA-sequencing reveals loss of metabolic liver zonation in O-GlcNAcylation deficient livers.
- Loss of O-GlcNAcylation promoted DEN-Induced HCC.
- Increase of hepatic O-GlcNAcylation prevented HCC progression.

## Introduction

O-GlcNAcylation is a dynamic post-translational modification (PTM) that involves the addition of N-acetylglucosamine (GlcNAc) onto proteins via an oxygen-linked bond. This process is regulated by two enzymes: O-GlcNAc transferase (OGT), which adds GlcNAc to proteins using UDP-GlcNAc as the GlcNAc donor, and O-GlcNAcase (OGA), which removes this modification (1). Because the hexosamine biosynthetic pathway, which produces UDP-GlcNAc, integrates multiple metabolic pathways, including nucleotide, fatty acid, protein, and glucose metabolism (1), changes in multiple independent metabolic pathways can affect O-GlcNAcylation. Previous studies show that O-GlcNAcylation plays a role in a number of cellular processes, including metabolism, inflammation, and cell proliferation (1, 2). Abnormal O-GlcNAcylation levels have been linked to various diseases, including cancer (1, 3, 4). A recent study by our group found that a lack of hepatic O-GlcNAcylation during liver regeneration impairs the termination phase and leads to sustained cell proliferation and loss of hepatocyte identity (2). Whereas O-GlcNAcylation is known to be involved in hepatic fibrosis and HCC pathogenesis, the mechanisms are not clear (4-6). These findings suggest that maintaining proper levels of O-GlcNAcylation may be important for the maintenance of healthy cells and the prevention of certain degenerative diseases.

During disease progression in the liver, hepatocyte nuclear factor 4 alpha (HNF4α), a critical regulator of maintaining hepatocyte differentiation, is known to decrease, causing a decrease in liver function and hepatocyte dedifferentiation (7, 8). This leads to increase cell proliferation ultimately increase susceptibility to progression to HCC (7, 8). O-GlcNAcylation has been shown to be a critical regulator in cellular differentiation, such as hematopoietic stem cells and neuronal cells (9, 10). Previous studies from our lab have shown cross-talk between O-GlcNAcylation and HNF4α (2). Here, we studied the effects of chronic loss of O-GlcNAcylation and the role of O-GlcNAcylation in the progression of liver disease. Using hepatocyte-specific OGT knockout mice and single-cell RNA-sequencing technologies, we found that loss of O-GlcNAcylation leads to the loss of liver zonation, hyperplasic hepatocyte nodules, and significant dedifferentiation. Further, we identified multiple molecular mechanisms at play in regulation of hepatic differentiation by O-GlcNAcylation. Our data suggest that O-GlcNAcylation is a potential therapeutic target in the maintenance of hepatic differentiation during the progression of liver disease.

## Methods

### Human Liver Samples

Human tissues were obtained from the KUMC Liver Center (normal n of 4, NASH n of 3, NASH + cirrhosis n of 4). All human liver tissues were obtained with informed consent in accordance with ethical and institutional guidelines. All studies were approved by the Institutional Review Board of KUMC. HCC human tissue microarray (TMA) was purchased from US Biolab (cat # LIV048-10A). The human protein atlas tool was utilized to visualize OGT protein levels in human HCC samples (TMA #T-56000) (11).

### Animal Care and Models

The Institutional Animal Care and Use Committee (IACUC) at the University of Kansas Medical Center approved all animal studies and was performed in accordance with IACUC regulations. All mice were housed in the KUMC vivarium with a standard 12-hour light and 12-hour dark cycle. Generation of genetically altered OGT-floxed mice and OGA-floxed has been previously described (12, 13). Both OGT-floxed and OGA-floxed were bred on a C57BL/6J background. Hepatocyte-specific knockouts were generated by injecting AAV8-TBG-CRE and using AAV8-TBG-eGFP (Vector Biolabs) as a control, as previously described (2).

For chronic OGT deletion, two-month-old male OGT-floxed mice were injected intraperitoneally (i.p.) with AAV8-TBG-GFP or AAV8-TBG-CRE, and tissues were collected 35 days after AAV8 administration. To study liver cancer pathogenesis, OGT-floxed and OGA-floxed pups were treated with diethylnitrosamine (DEN,15 ug/kg i.p.) on postnatal day 15. Five months after DEN injections, floxed mice were injected with AAV8-TBG-Cre to knockout OGT or OGA. Controls were generated by injecting AAV8-TBG-EGFP. Mice were euthanized two months after AAV8 injection. In all studies, liver injury was assessed by serum ALT activity assay, and livers were weighed after cholecystectomy to calculate liver-weight-to-body-weight ratios as previously described (2). Glucose was measured in serum according to the manufacturer’s protocol (Pointe Scientific cat# G7521120).

### Staining Procedures and Imaging

Paraffin-embedded liver sections (5 μm thick) were used for hematoxylin and eosin staining (H&E), immunohistochemistry, and picrosirius red staining, as previously described (2, 14). Primary and secondary antibodies with respective dilutions are shown in **Table S1**. Photomicrographs were captured with an Olympus DP74 color camera mounted on an Olympus BX51 microscope with CellSens (Version 2.3) software.

### Protein Isolation and Western Blotting

Approximately 100 mg of liver tissue was homogenized using a beaded tube containing 300 μL of RIPA buffer (20 mM pH 7.5 Tris, 150 mM NaCl, 2 mM EDTA, 1 mM DTT, 40 mM GlcNAc, 0.1% Sodium Deoxycholate Acid, 0.1% SDS, and 1%NP-40) containing 1x Halt Phosphatase inhibitor and protease cocktail (Thermofisher cat# 78427 & 78438). Pierce BCA Protein Assay (Thermofisher cat# 23225) was used to measure protein levels as previously described (15). Antibodies used with respective dilutions are shown in **Table S1**. All western blots were imaged with either SuperSignal West Pico PLUS or Femto Maximum (Thermofisher cat# 34578 and 34096). Western blots were imaged on Odyssey LiCor utilizing Image Studio software (Version 5.2).

### RNA Isolation and qPCR

RNA was isolated from ∼25 mg of liver tissue utilizing the TRIzol method according to the manufacturing protocol (Thermofisher cat# 15596026). Isolated RNA concentration was measured using an Implen N50 Nanophotometer. cDNA was made with 2000 ng of RNA per reaction utilizing the High-Capacity cDNA Reverse Transcription Kit according to the manufacturer’s protocol (Thermofisher cat# 4368814). qPCR was performed using 50 ng per reaction with PowerUp SYBR green master mix and final concentration of 2.5 μM of forward and reverse primers (**Table S2**) according to the manufacturer’s protocol (Thermofisher cat# A25741). A BioRad CFX384 system was used to run qPCR reactions in a 384-well plate setup. Raw data were analyzed using the CFX Maestro 1.1 Software (Version 4.1.2433.1219).

### Single-Cell Suspensions

Livers of chronic deleted OGT-KO mice (30 days) and controls were perfused utilizing two-step collagenase perfusion to isolate parenchymal and nonparenchymal cells (NCPs), as previously described (16). Trypan blue and a hemocytometer were utilized to determine cell viability and concentration of cells. All samples yielded a viability greater than 85%. A 50g centrifugation was used to pellet parenchymal cells from NPCs. Percoll gradients were used to remove dead cells. 100% buffered Percoll was made by a 1:9 ratio of 10x PBS to Percoll (Fisher Scientific cat# 45-001-747).

Parenchymal cells were resuspended in 50% buffered Percoll and 50% isolation media (Dulbecco’s Modified Eagle Medium (Corning™ cat# 10014CV) supplemented with 10% fetal bovine serum (FBS) and 2% bovine serum albumin (BSA)) and centrifuged at 72g to remove dead cells. The parenchymal pellet was resuspended in isolation media to reach a concentration of 500 cells/μL. NPCs were pelleted using 600g centrifugation. 3 mL of red blood cell (RBC) lysis buffer (155 mM ammonium chloride, 10 mM potassium bicarbonate, 0.1 mM EDTA, pH 7.3) was added to remove RBC contamination for 3 minutes. NPCs were then resuspended to a final concentration of 35.3% buffered Percoll and 64.7% isolation media and centrifuged at 900g for 10 minutes. The NPC pellet was then resuspended in isolation media for a concentration of 1200 cells/μL.

### Single-Cell RNA-Sequencing

Single-cell suspension of hepatocytes and NPCs from OGT-KO and control mice was used for single-cell RNA-sequencing using the 10× Chromium Single Cell 3′ Gene Expression profiling platform targeting approximately 10 000 cells per sample and a read depth of 50 000 read per cell, as previously described in depth (17). Single-cell libraries were then sequenced using the Illumina NovaSeq 6000 S1 Flow Cell for 100□cycles. Raw data were analyzed utilizing 10x Cell Ranger using the mkfastq pipeline (Version 6.0.2) and aligned to the mouse transcriptome mm10-2020-A using the Apte Lab server (HPE DL380 Gen10 8SFF CTO high-performance server) (18). Raw data were deposited into the GEO database (GSE223830). The aligned barcodes, features, and matrix files were uploaded into RStudio (Version 4.0.3, RStudio Team).

### Data Analysis of Single-Cell RNA-Sequencing

Data analysis was performed using the Seurat Package (Version 4.0.3). Single-cell data from parenchymal cells and NPCs for OGT-KO mice and controls were cleaned independently. For parenchymal cell analysis, duplicate cells or cells without a low read count and a high percentage of mitochondrial genes were kept in the analysis (if the number of captured features was between 100 and 7000, with mitochondrial genes being < 75% of the features). The cells were then log normalized, scaled using a linear model, utilized the first 20 PCA dimensions to determine nearest neighbors, and a resolution of 0.2 to generate clusters. Hepatocyte markers were identified utilizing previously established markers, and all other cell types were filtered out (17). For the NPCs, cells were filtered using features between 100 and 7000, with mitochondrial genes being < 20% then used a resolution of 0.4 for clustering. The markers within the Immunological Genome Project (ImmunoGen) database were used to identify populations within the NPC fraction. Hepatocytes were then filtered out of the NPCs. After data clean-up, the control NPCs were merged with OGT-KO NPCs and control hepatocytes were merged with OGT-KO hepatocytes to produce an NPC and hepatocyte object. These two objects were then clustered using 0.2 resolutions for both the NPC and hepatocytes. The cell types were then further annotated, including genotype labels. Finally, the merged hepatocyte and NPC objects were merged and clustered using a 0.2 resolution. Cell contamination was exhibited between some clusters, likely due to a technical error, with cells clumping upstream of the sequencing. Each cell type was then subclustered to remove any residual contamination and reclustered to determine fine cell type labels using the ImmunoGen database. To predict cell-cell communication, we utilized the package CellChat (Version 1.5.0) (19). The package standard workflow was utilized using the mouse interaction database.

### RNA-Seq Data Acquisition

An online OGT dataset in rodents was downloaded using SRA-tools from the GEO database (GSE188882) (2). Raw fastq files were then aligned to the mouse genome (GRCm38) and counted using STAR software (20). DESeq2 (Version 1.28.1) in R Studio (Version 4.0.3, RStudio Team) was used for count normalization and differentially expressed gene (DEG) lists, as previously described (7).

### The Cancer Genome Atlas

The RStudio (Version 4.0.3, RStudio Team) package TCGAbiolinks (Version 2.16.4) was utilized to download liver hepatocellular carcinoma (TCGA-LIHC) RNA-sequencing data (21). The prebuilt EdgeR pipeline was utilized to generate DEG lists with a log transformation. Analyzed data were annotated with biomaRT (Version 2.44.4) using the Ensembl database.

### Statistical Analysis and Data Visualization

For experiments not associated with single-cell or bulk RNA-seq, such as ALT measurements, the results are expressed as mean ± standard error of the mean (SEM). Bar graphing and statistical analysis were carried out in GraphPad Prism 9. Student’s two-tail t-test or two-way ANOVA with Sidak’s post-hoc test was applied to all analyses, with a p-value <0.05 considered significant. Dot plots, heatmaps, Venn diagrams, and UMAPs were produced in RStudio (R version 4.0.3; RStudio Team).

## Results

### Hepatic O-GlcNAcylation levels decrease in chronic liver disease progression in humans

Western blot analysis using human liver samples of normal, steatosis, nonalcohol steatohepatitis (NASH), cirrhosis, and HCC showed a progressive decline in total O-GlcNAcylation (**Fig. 1A**). Normal livers had the highest total hepatic O-GlcNAcylation, whereas cirrhosis and HCC had the lowest. There was no difference in OGT protein levels in healthy, steatosis, and NASH samples, but OGT was completely absent in cirrhosis and HCC samples. IHC of O-GlcNAcylation in human liver samples of NASH or NASH+cirrhosis corroborated that O-GlcNAcylation is maintained in NASH without cirrhosis but is decreased in NASH with cirrhosis (**Fig. 1B)**. Immunohistochemistry (IHC) of O-GlcNAcylation on tissue microarrays containing normal and HCC samples showed decreased O-GlcNAcylation and OGT in HCC (**Fig. 1C-D**). Taken together, these data indicate that O-GlcNAcylation is lost during late-stage liver disease.

**Figure 1.**
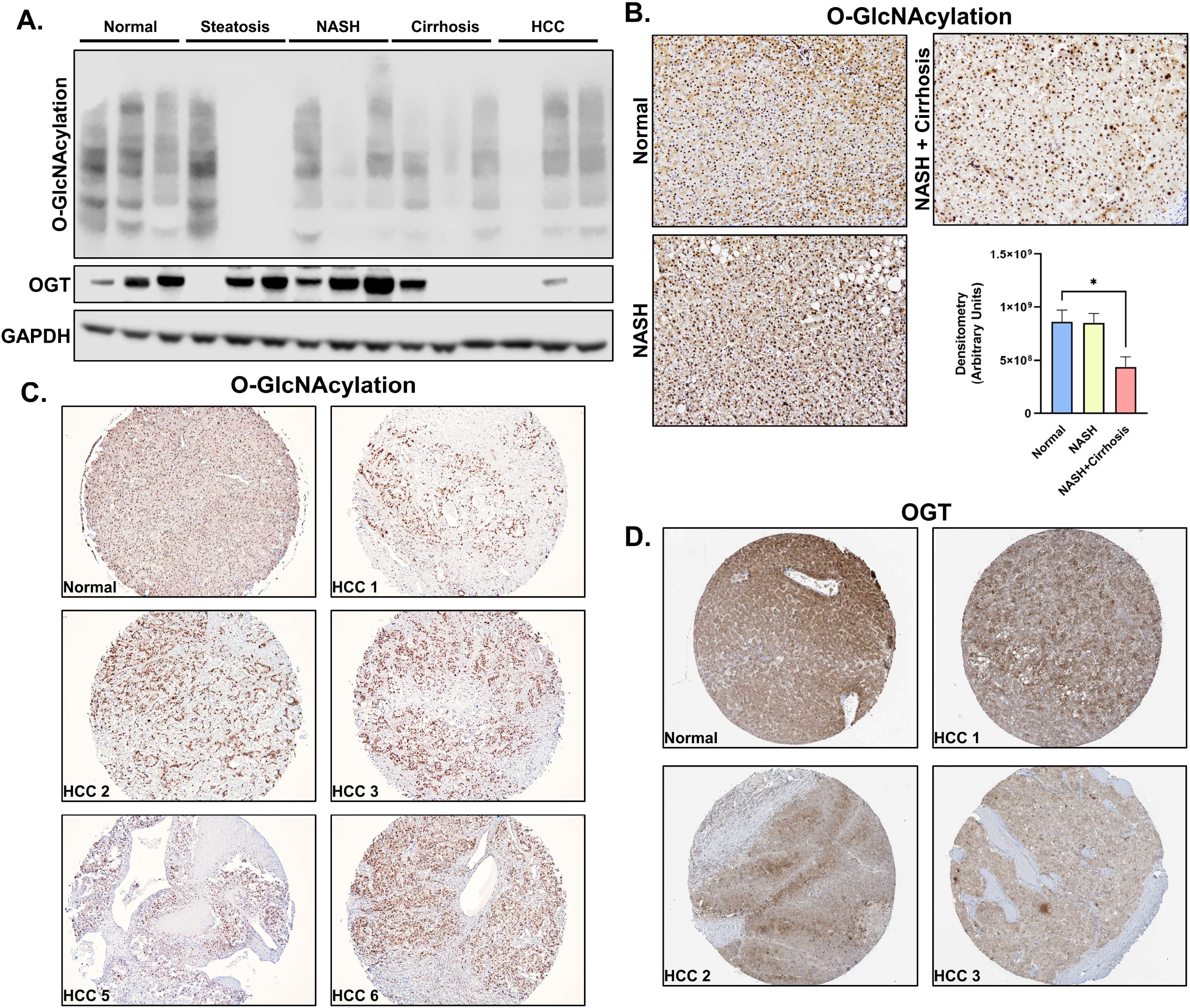
O-GlcNAcylation is lost during the progression of liver disease. (A) Western blot analysis of human liver samples for total O-GlcNAcylation, OGT, and the housekeeping protein GAPDH. (B) IHC of total O-GlcNAcylation from liver tissue microarrays of human hepatocellular carcinoma samples and control livers. IHC of (C) O-GlcNAcylation of HCC and normal human samples derived from TMA and (D) OGT derived from the human protein atlas.(E) Heatmap of fold changes of common genes derived from RNA sequencing from livers of OGT-KO mice compared to controls after PHX (GSE188882) and HCC compared to healthy human tissues (TCGA-LIHC). Orange and blue represent positive and negative log_2_(Fold Change), respectively. (D) IHC of total O-GlcNAcylation of livers from healthy, NASH, and NASH with cirrhosis human liver tissue.

### O-GlcNAcylation is required to maintain liver homeostasis and contributes to the maintenance of hepatic zonation

To study the effects of a chronic loss of O-GlcNAcylation in hepatocytes, we studied histopathological and molecular changes in hepatocyte-specific OGT-KO mice 35 days after OGT deletion. Western blot analysis was utilized to confirm successful OGT-KO (**Fig. 2B**). OGT-KO mice exhibited significant hepatomegaly, as indicated by an increase in liver-weight-to-body-weight ratio, and increased liver injury (**Fig. 2C-D**). H&E staining of OGT-KO livers showed significant hepatocyte dysplasia (**Fig. 2E**), which was accompanied by reorganization of F4/80^+^ macrophages (**Fig. 2F**), a significant increase in αSMA, a marker for activated hepatic stellate cells (HSCs) (**Fig. 2G**). Picrosirius red (PSR) staining revealed significant collagen deposition in OGT-KO livers (**Fig. 2H**).

**Figure 2.**
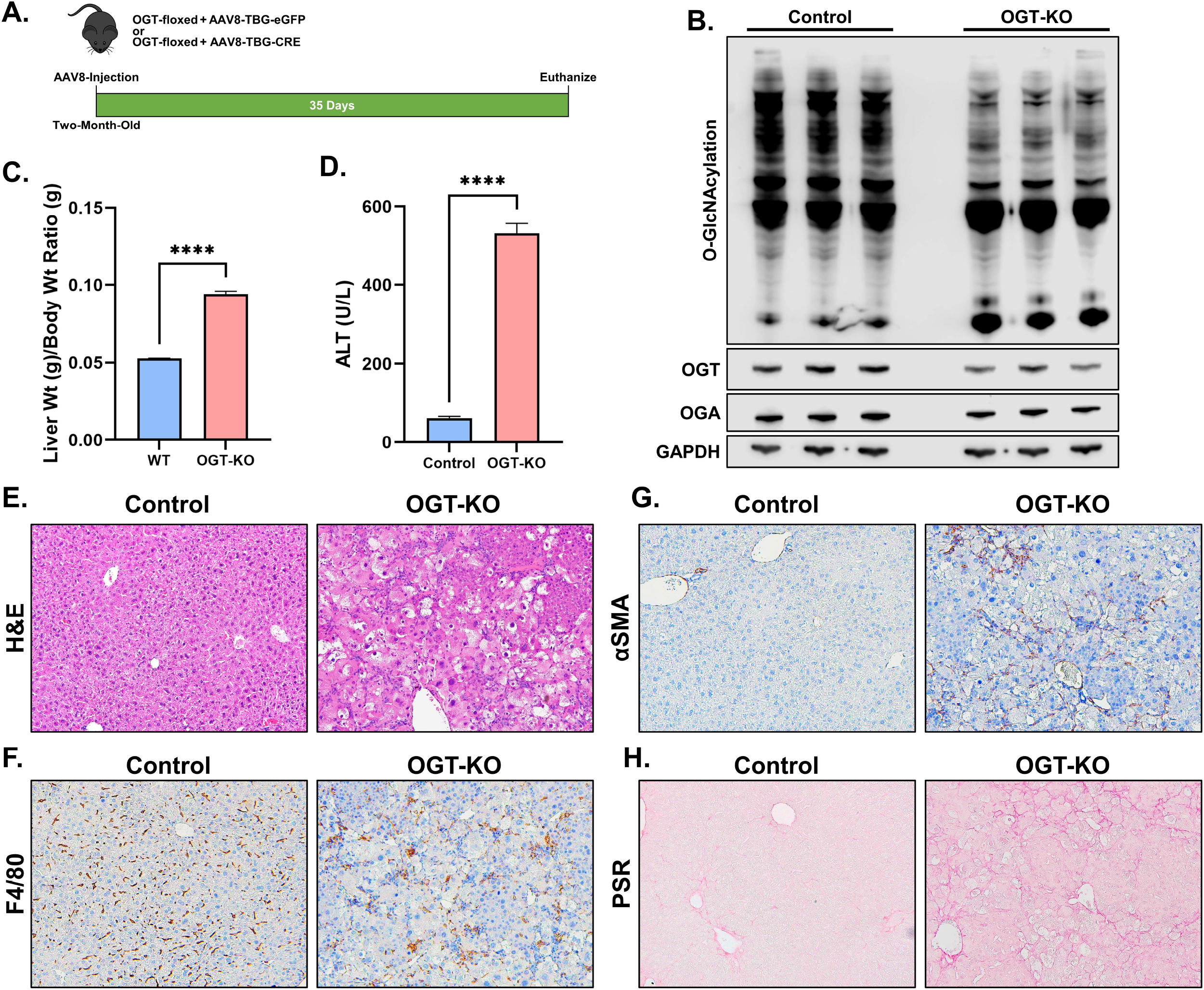
Chronic deletion of OGT leads to dysplastic liver lobules. (A) Western blot analysis of hepatic O-GlcNAcylation, OGT, OGA, and GAPDH from 35-day OGT-KO mice. (C) Liver weight-to-body weight ratios and (D) ALT serum levels of control and OGT-KO mice. Bar represents the mean, with error bars representing SEM. (E) H&E microsections of control and OGT-KO mice 35 days after deletion. Immunohistochemistry of (F) the macrophage marker F4/80 and (G) the activated hepatic stellate cell marker αSMA in control and OGT-KO mice (200x magnification). (H) Picrosirius red (PSR) staining to visualize collagen deposition in control and OGT-KO mice (200x magnification). Level of significance: ****p < 0.0001; **p < 0.01 (Two-tailed t-test)

To interrogate changes in specific cell populations, we turned to single-cell RNA-sequencing (scRNA-seq). Unsupervised cell clustering produced 11 unique clusters identified as Kupffer (Kup) Cells, Hepatocytes (Heps), Endothelial (Endo) Cells, B-cells, Natural Killer (NK) Cells, T-cells, Dendritic Cells (DC), Neutrophils (Neut), and Monocytes (Monos) in both control and OGT-KO mice (**Fig. S1A-C**). Specific markers were used to identify each cluster (**Fig. S1D**). T-cells and hepatocytes were the most captured cells in the control and OGT-KO mice, respectively. The least common were monocytes (control) and endothelial cells (OGT-KO) (**Fig. S1E-F**). HSCs were not captured during the single-cell isolation.

The hepatocytes exhibited the greatest difference at a single-cell resolution. The OGT-KO hepatocytes exhibited minimal overlap with the control hepatocytes (**Fig. 3A**). We subclustered the hepatocytes into either periportal (PP), midzonal (MZ), or perivenous hepatocytes (PV) using following gene markers: *Glul, Lect2*, and *Cyp2e1* for PV; *Alb, Cdh1*, and *Cyp2f2* for PP hepatocytes. The control liver lobule had distinct PP and PV hepatocyte populations, with MZ having an overlap of both PP and PV markers, particularly *Cyp2e1* and *Cyp2f2* (**Fig. 3B-C**). The chronic deletion of OGT showed a complete loss of PP features (**Fig. 3C**). An additional population that lacked both PP and PV features was identified in the OGT-KO mice, which we termed as dedifferentiated (DF) hepatocytes. We confirmed the scRNA-seq data by performing IHC for CYP2F2 and CYP2E1 on control and OGT-KO liver sections. Consistent with scRNA-seq data, the control livers showed strong staining of CYP2F2 in the PP region and CYP2E1 in the PV region (**Fig. 3D**), which was significantly lower in OGT-KO hepatocytes (**Fig. 3D**).

**Figure 3.**
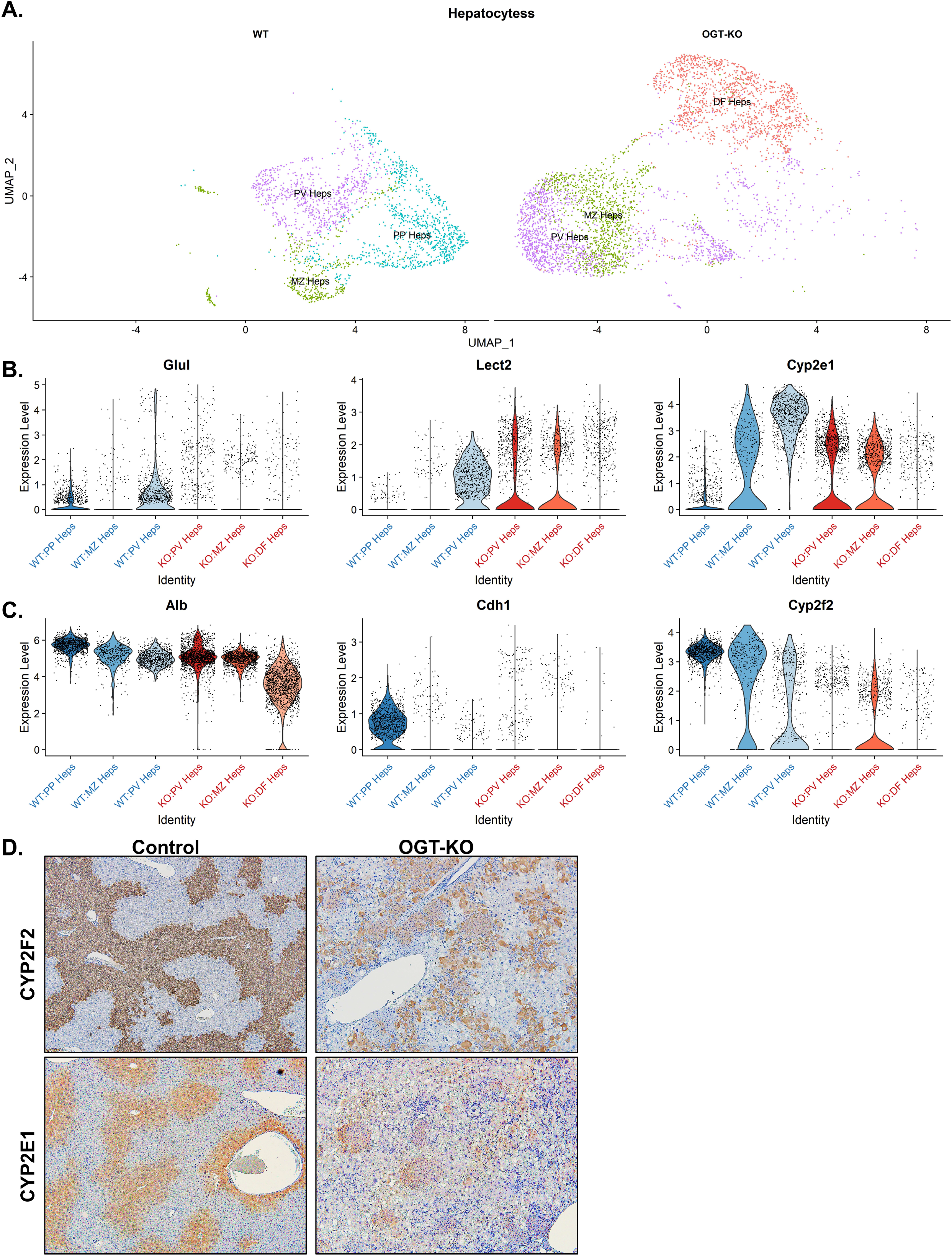
Single-cell RNA sequencing analysis of hepatocytes derived from 35-day OGT-KO mice. (A) UMAP plot of SC RNA-seq of hepatocytes split by control (WT) and 35-day OGT-KO mice. Violin plots of (B) perivenous (PV) markers (*Glul, Lect2*, and *Cyp2e1*) and (C) periportal (PP) markers (*Alb, Cdh1*, and *Cyp2f2*). Each black dot represents a hepatocyte with the distribution of the expression levels. (D) Immunohistochemistry of CYP2F2 and CYP2E1 in control and OGT-KO livers (200x magnification).

Previous studies have shown that loss of O-GlcNAcylation results in increased NRF2 activity (22, 23). Consistent with these findings, the hepatocytes in the OGT-KO mice exhibited increased expression of the target genes *Gsta2, Gstm1, Txn1*, and *Gclc*, especially in the DF population (**Fig. 4A**). To further characterize the DF population, we generated three differentially gene lists of the comparisons KO:DF vs WT:PP, vs WT:PV, and vs WT:MZ. Pathway analysis using Metascape was performed on these gene lists. HNF4α and HNF1α was the most significantly impacted regulators (**Fig. 4B**). Target genes of HNF4α were accessed in all hepatocyte populations and found decreased expression in *Apob, Apoa2, Cyp227a1, Dio1, Pck1*, and *Ugt2b1* in the OGT-KO populations compared to control. Notably, the DF Heps were the most affected compared to the OGT-KO MZ and PV populations (**Fig. 4C**). FOXO1 was another impacted factor that is known to co-regulate gene expression with HNF4α (24) and is important in metabolic zonation of the liver, particularly glucose metabolism. To investigate this further, FOXO1 target genes were interrogated in the scRNA-seq dataset. Interestingly, all three populations in the OGT-KO hepatocytes showed loss of FOXO1 activity (**Fig. 4D**). To interrogate other methods of glucose metabolism, we performed Periodic acid–Schiff (PAS) staining on control and OGT-KO mice. As expected, glycogen was concentrated in the PP region in the control mice, whereas in the OGT-KO mice, glycogen storage did not show zonation, and a number of hepatocytes exhibited complete loss of glycogen throughout the liver lobule (**Fig. 4E**).

**Figure 4.**
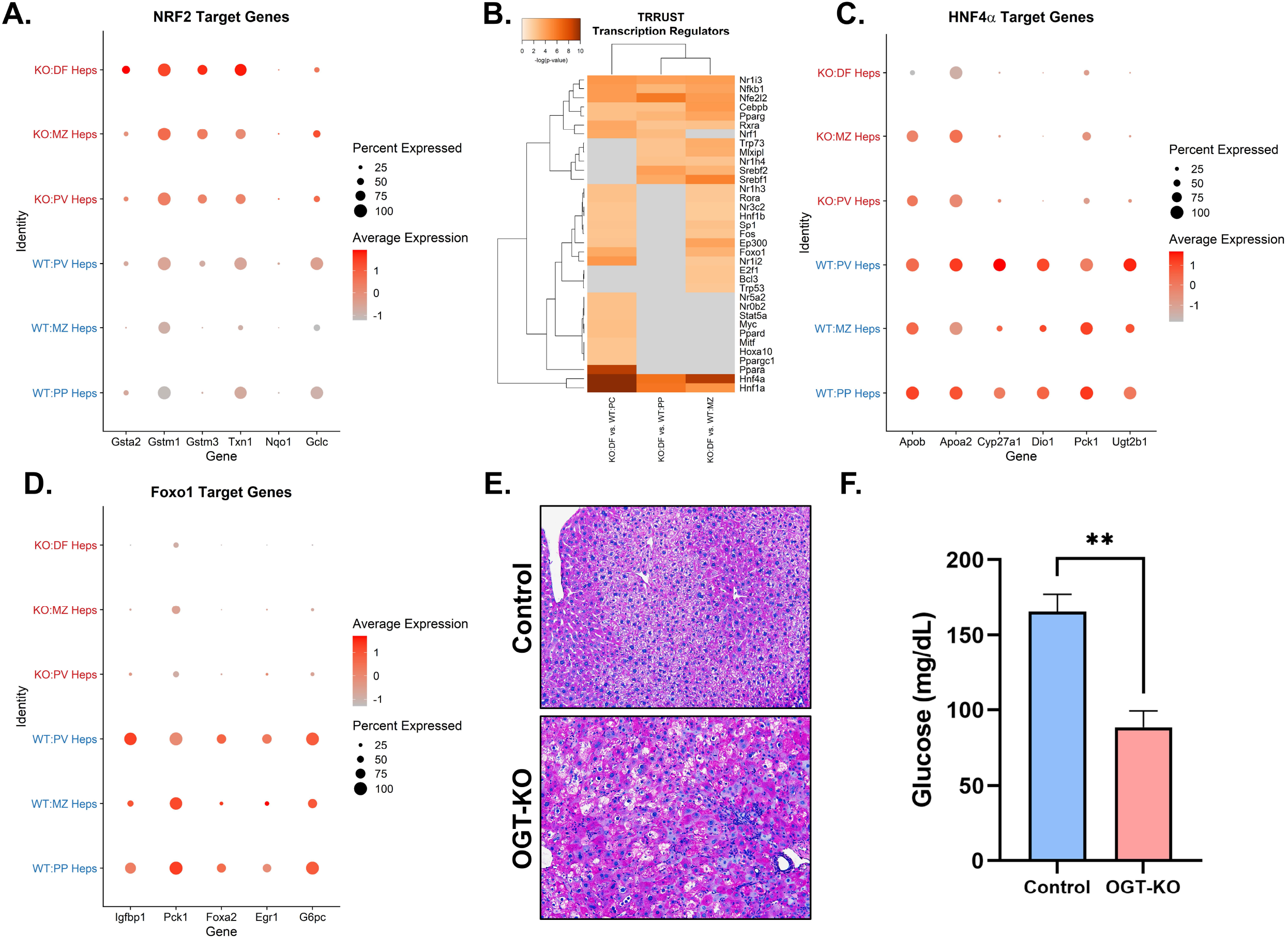
OGT-KO mice had decreased HNF4α activity and altered metabolic zonation of glycogen storage. (A) Dot plot of NRF2 target genes in each population represented in a dot plot. (B) Heatmap of the -log(pvalue) for the altered TRRUST transcription regulators comparing the dedifferentiated hepatocytes to each control population. Gray represents no assigned p-value, and orange scale represents -log(pvalue). Dot plot of (C) HNF4α and (D) FOXO1 target genes in each population, represented in a dot plot. (E) Periodic Schiff staining of livers of control and OGT-KO mice to visualize glycogen (200x magnification). In the dot plots, the size represents the percentage of cells that express that gene, and color represents the average expression in the cell population. (F) Bar graph of serum glucose levels in the control and OGT-KO mice. Bar represents the mean, with error bars representing SEM. Level of significance: **p < 0.01 (Two-tailed t-test)

To determine whether excess storage was due to hyperglycemia, we measured the glucose levels in the serum and found a significant decrease in the OGT-KO, indicating a flux of serum glucose to glycogen in the liver (**Fig. 4F**). These data indicate that O-GlcNAcylation is critical for maintaining liver lobule zonation through HNF4α.

The NPC populations were then subclustered to determine changes in the OGT-KO (**Fig. 5A-E**). The most striking differences of the NPCs were increasing infiltrating monocytes and Kupffer cells (**Fig. 5B**), the change in the CD4^+^ T-cells population and increased of CD8^+^ T-cells (**Fig. 5C)**, and an increase in Fcer2a^+^ B cells (**Fig. 5D**). Cell-cell communication analysis of the fine-labeled populations showed that the main signal transponders were endothelial cells and the main signal receivers were CD8^+^ T-cells (**Fig. 5F-G**). This indicates that endothelial cells could play a role in T-cell recruitment in the OGT-KO livers.

**Figure 5.**
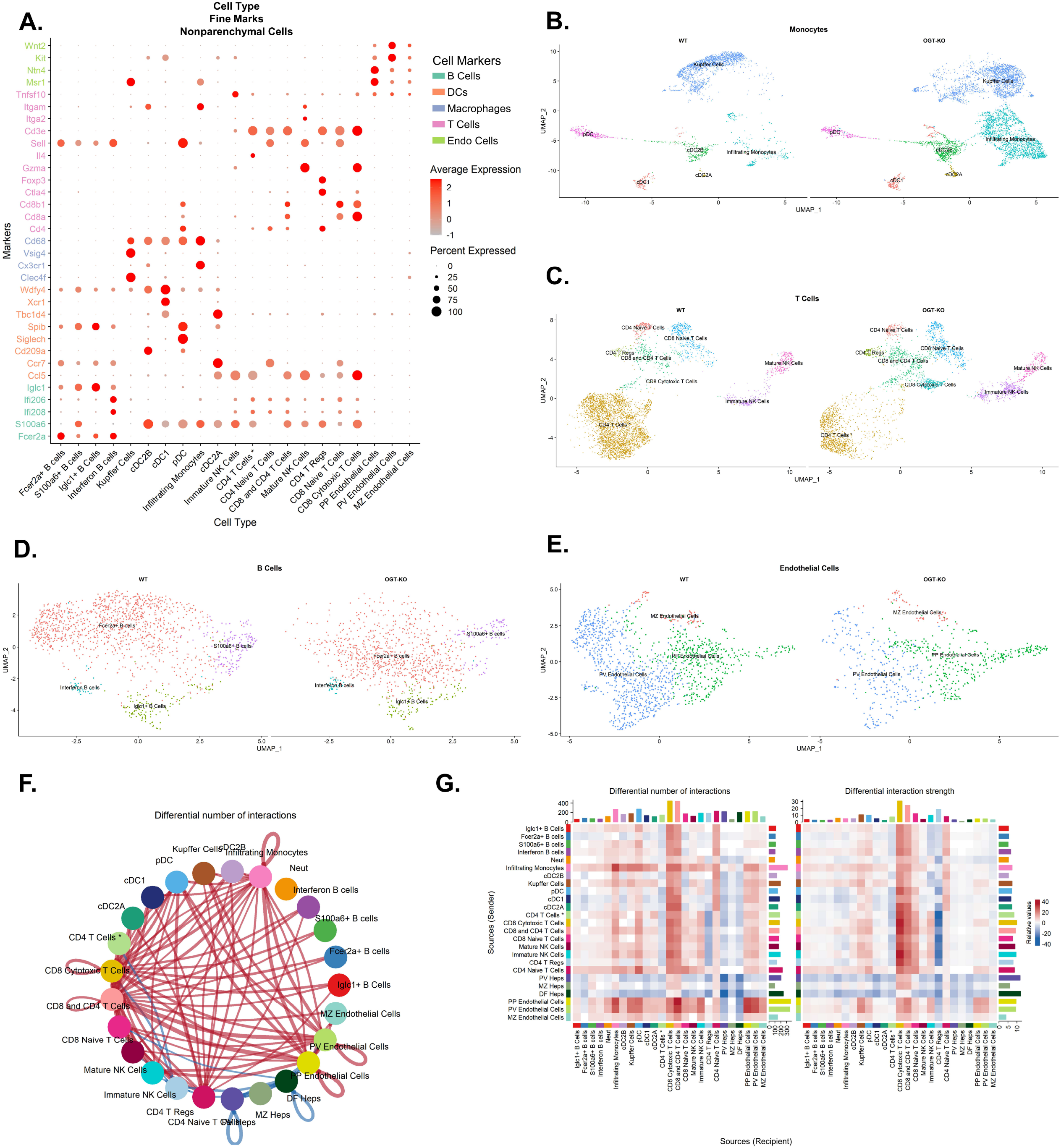
OGT-KO NPC populations significantly changed in the OGT-KO mice. (A) Dot plot of fine cell-type markers. The color of the dot represents the expression level, and the size of the dot represents the percentage of cells expressing the marker. Gene name color represents the larger category of cell types. UMAP of subcluster with fine labels of (B) monocytes, (C) t-cells, (D) B-cells, and (E) endothelial cells. (F) Cord diagram of predictive interaction between populations defined by fine cell type. Red and blue connections represent increased and decreased signaling in OGT-KO, respectively. Size of the connection indicates the number of cell-cell interactions. (G) Heatmap of differential number of interactions and interaction strength. Color scale represents increased (red) and decreased (blue) signals in OGT-KO mice compared to the control. Side bar graphs represent the sum of outgoing signals for the number and strength of interactions per cell type. The top columns represent the sum of incoming signals for the number and strength of interactions per cell type.

### Depletion of O-GlcNAcylation promotes diethylnitrosamine-induced HCC

Next, we investigated if loss of O-GlcNAcylation promotes the development of HCC using the diethylnitrosamine (DEN)-induced HCC model (25). DEN was injected into OGT-floxed mice 15 days postnatal to initiate HCC development. OGT was deleted using the AAV8 system 5 months after DEN injection, and 2 months were allowed for tumor promotion (**Fig. 6A**). Western blot analysis confirmed a decrease in total O-GlcNAcylation and OGT in OGT-KO mice compared to their control groups (**Fig. 6B**). Importantly, OGT-KO mice had significantly more tumors compared to the control group (**Fig. 6C-D**), with an increased liver-weight-to-body-wight ratio and liver injury (**Fig. 6E-F**). H&E staining showed OGT-KO livers had significant neoplasia (**Fig. 6G**), which was deemed to be HCC based on markers such as CK8, reticulin, and glypican 3 (**Fig. 6H**). Because our studies show that O-GlcNAcylation is required for hepatic differentiation, we determined if there is increased stemness in OGT-KO livers with tumors. qPCR analysis showed significant increase in stemness markers, including *Nanog, Klf4, Myc*, Sox2, and *Pou5f1* (OCT4 gene), in OGT-KO livers. During chronic liver disease HNF4α function is known to decrease (7). We measured expression of genes regulated either positively (*Ugt2b1, Dio1, Apoa2*, and *Ces3*) or negatively (*Ect2* and *Akr1b7*) by HNF4α and found significant decrease and increase, respectively (**Fig. 6J-K**).

**Figure 6.**
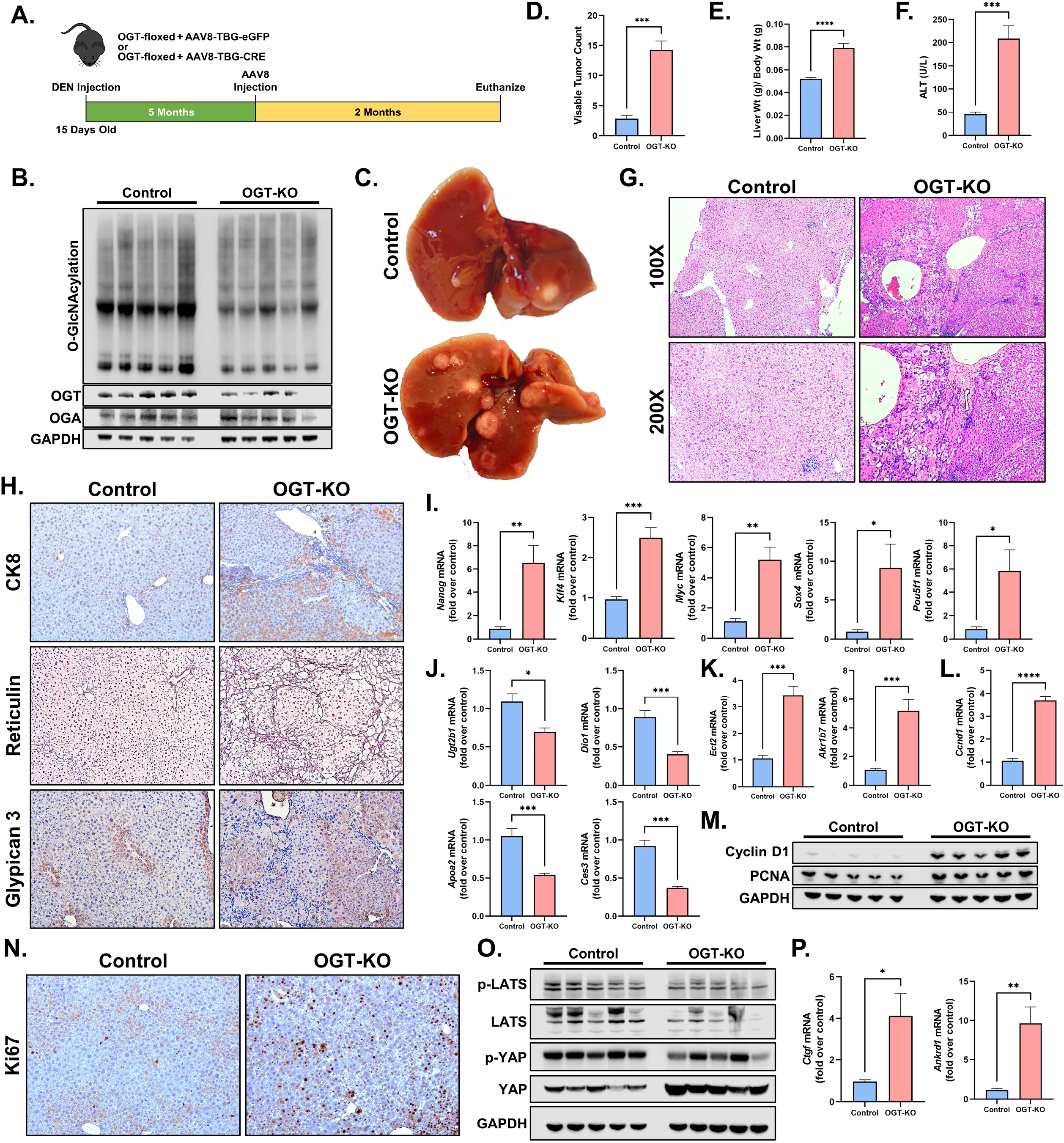
O-GlcNAcylation is an impediment to the progression of DEN-induced HCC. (A) Experimental design of DEN-induced HCC. (B) Western blot analysis of hepatic O-GlcNAcylation, OGT, OGT, and GAPDH in control and OGT-KO mice. (C) Gross photos of livers from control and OGT-KO after DEN-induced HCC. Bar graphs of (D) visible tumor count, (E) liver-weight-to-body-weight ratio, and (F) serum ALT of control and OGT-KO mice. (G) Photomicrographs of H&E of control and OGT-KO at 100x and 200x magnifications. (H) Tumor markers, including CK8, Reticulin, and Glypican 3, in control and OGT-KO mice (200x magnification). qPCR of genes that regulate (I) hepatocyte stemness and HNF4α (M) upregulated and (N) downregulated target genes. (L) qPCR of cyclin D1, and (M) western blot analysis of the cell proliferation markers cyclin D1 and PCNA. (N) Immunohistochemistry of the proliferative marker Ki67 (200x magnification). (O) Western blot analysis of proteins involved in YAP signaling pathway. (P) qPCR of YAP target genes. For qPCR, values were normalized to 18s then to the control group. Bars represent mean, with error bars representing the SEM. Levels of significance: ****p < 0.0001; ***p < 0.001; **p < 0.01; *p < 0.05 (Two-tailed t-test)

To determine the extent of cell proliferation, we measured the expression of cyclin D1 and found that it was significantly upregulated in OGT-KO mice (**Fig. 6L**). This was corroborated by increased protein levels of cyclin D1, with a slight increase in PCNA in OGT-KO mice (**Fig. 6M**). IHC of Ki67 showed an increase in hepatocyte proliferation, as well as NPC proliferation (**Fig. 6N**). To investigate the mechanisms of increased DEN-induced carcinogenesis in OGT-KO mice, we investigated WNT, AKT, and ERK signaling, and the Hippo Kinase pathway, all of which is known to be activated in HCC. Western blot analysis showed a significant decrease in ERK and AKT activity (**Fig. S2A-B**). No changes were exhibited in either phosphorylated ß-catenin (inactive) or unphosphorylated ß-catenin (active) (**Fig. S2C**). qPCR of ß-catenin target genes showed no changes in *Axin2* and a significant suppression of *Cyp2e1* and *Glul* (**Fig. S2D**). Lastly, we found decrease in phosphorylated LATS and phosphorylated Yap but an increase in total Yap in OGT-KO mice compared to control mice (**Fig. 6O**). qPCR on YAP target genes (*Ctgf* and *Ankrd1*) corroborated the YAP activity data (**Fig. 6P**).

These data indicated that proliferation is governed by YAP signaling. OGT-KO mice exhibited a significant induction in the proinflammatory markers *Adgre1, Tnfa*, and *Il6*, inflammatory nodules, and NFκB signaling (**Fig. 7A-C**). qPCR on profibrotic genes (*Tfgb1, Des, Acta2, Col1a1, Col1a2*, and *Col1a3*) was performed and found a significant induction in OGT-KO mice, which was corroborated by αSMA IHC and PSR staining (**Fig. 7D-F**).

**Figure 7.**
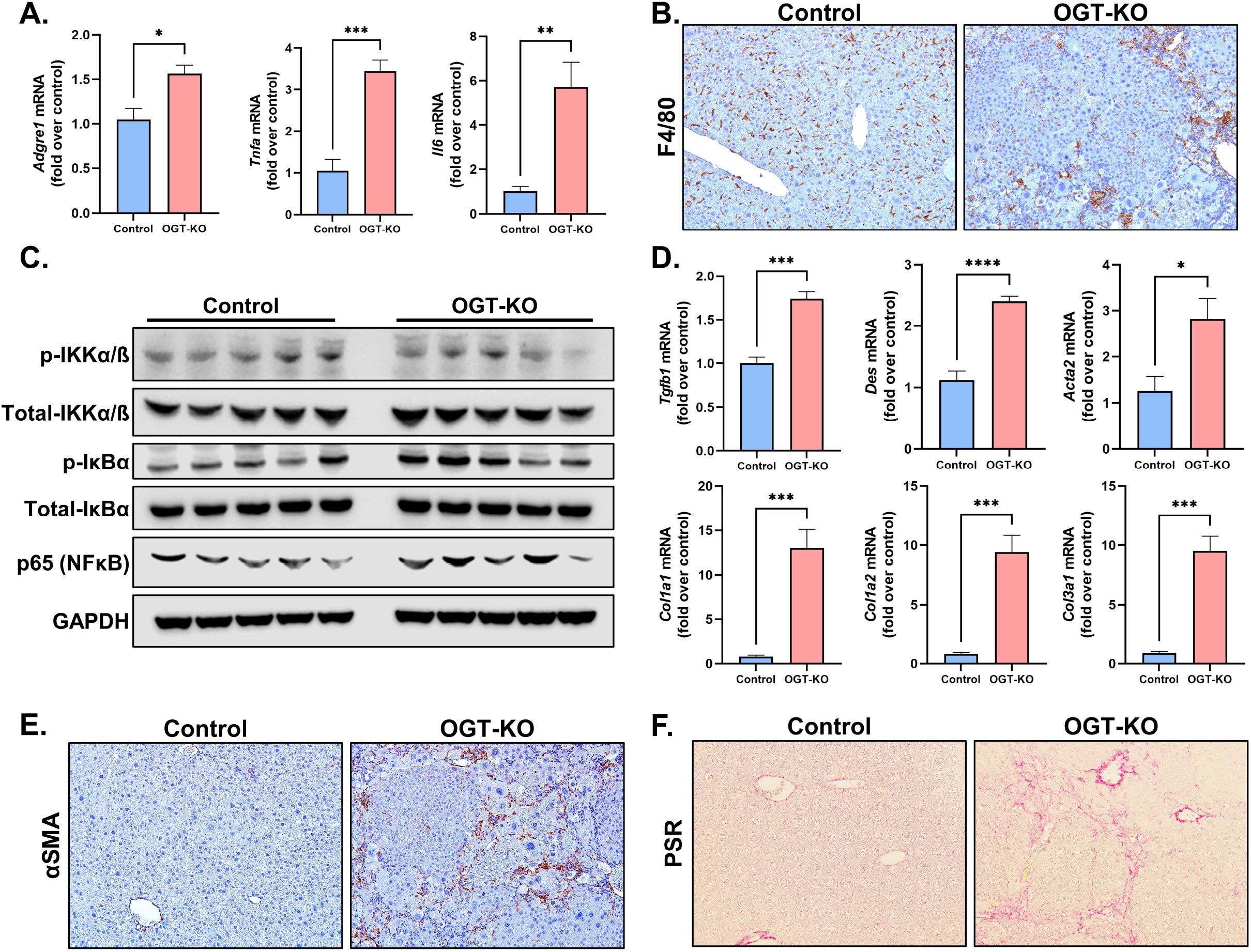
OGT-KO exhibited increased inflammation and fibrosis during the promotion of DEN-induced HCC. (A) qPCR of proinflammatory markers (*Adgre1, Tnfa*, and *Il6*). (B) Immunohistochemistry of F4/80. (C) Western blot analysis of NFkB pathway including phosphorylated /total Ikka/b, phosphorylated/total IkkBα, p65, and GAPDH. (D) qPCR of profibrotic genes (*Tgfb1, Des, Acta2, Col1a1, Col1a2*, and *Col3a1*). (E) Immunohistochemistry of αSMA and (F) picrosirius red staining in OGT-KO mice and controls. Photomicrographs are 200x magnification. Bars represent mean, with error bars indicating SEM. Level of significance: ****p < 0.0001; ***p < 0.001; **p < 0.01; *p < 0.05 (Two-tailed t-test)

To determine the effect of increased O-GlcNAcylation on HCC progression, we repeated the experiments in OGA-floxed mice (**Fig. S3A**). We observed a striking reduction in visible tumors in the DEN-treated OGA-KO mice. However, there was no significant difference in liver-weight-to-body ratio, liver injury, histological changes, HCC markers, or cell proliferation between OGA-KO and control mice treated with DEN (**Fig. S3B-M**). Additionally, OGA-KO mice did not have changes in inflammation or fibrosis (**Fig. S4A-D**).

## Discussion

O-GlcNAcylation is involved in a plethora of cellular processes, such as metabolism, proliferation, and cell differentiation (1). Because of its critical role in cellular functions, dysregulation of O-GlcNAcylation is known to be involved in diseased states, such as inflammation and fibrosis, both of which are hallmarks of hepatocellular carcinoma (HCC). In this study, we found that hepatocyte-specific OGT-KO resulted in a loss of hepatocyte differentiation and disruption in metabolic liver zonation. Further, we found that hepatocyte-specific loss of O-GlcNAcylation results in promotion of carcinogen-induced HCC, while OGA-KO mice, with higher hepatic O-GlcNAcylation, are protected from HCC development.

Previous studies have found that increased O-GlcNAcylation is a driver of HCC progression (6, 26-29). Because our data contradict the hypothesis that increased O-GlcNAcylation is a promoter of HCC, we repeated DEN-induced HCC in OGA-KO mice. Interestingly, we found that increased hepatic O-GlcNAcylation had fewer visible tumors compared to their respective controls, indicating protection against HCC progression.

However, no differences were exhibited in cell proliferation markers compared to the control group. The HCC model we implied was only a 7 month after DEN-injections with only 2 month of OGA deletion. This is relatively short to allow HCC to develop. We suspect that this was the reason the OGA-KO lacked significant changes cell proliferation markers. Future studies should be done to promote the expansion of HCC in OGA-KO and control mice and using longer timepoints to determine the extent of HCC progression by increasing O-GlcNAcylation (30).

Single-cell technologies have become more widely used to study the liver in different states (17, 31-33). It is well established that the liver is metabolically zonated, particularly hepatocytes, which shows distinct molecular patterns linked to their metabolic function (34). To classify these hepatocytes, markers for specific zones, either more perivenous (PV)- or periportal (PP)-like hepatocytes have been determined (17, 35). We utilized single-cell technologies to interrogate the effect of a chronic 5-week depletion of O-GlcNAcylation in hepatocytes. At a single-cell level, very few hepatocytes overlapped (between control and OGT-KO) using unsupervised clustering methods, indicating substantial transcriptome changes. This high-resolution sequencing data illustrate that not only O-GlcNAcylation affect hepatocyte differentiation, but it also affected PP hepatocytes at a greater magnitude compared to PV hepatocytes at the transcriptional level. This is likely attributed to the loss of hepatocyte nuclear factor 4 alpha (HNF4α) in OGT-KO hepatocyte populations. It is well established that loss of HNF4α leads to the dedifferentiation of hepatocytes into hepatoblast-like cells, allowing them to be in a more proliferative state (36). Past studies from our lab showed that, HNF4α levels need to decrease for hepatocytes to proliferate and chronic lack of HNF4α leads to liver disease progression (7, 36, 37). Interestingly, after 2/3^rd^ partial hepatectomy in OGT-KO mice, HNF4α levels are lost during liver regeneration, leading to a dedifferentiated phenotype (2). HNF4α is known to upregulate PP genes while suppressing PV features in PP populations (38). Conversely, in the PV region HNF4α is suppressed by LEF1, a WNT regulated gene, to prevent PP features. A functional example is that HNF4α and FOXO1 are known to coregulate glycogen metabolism, which is metabolically zonated (24). Our data show that glycogen storage is no longer localized in the PP region but exists throughout the liver lobule. These data illustrate that O-GlcNAcylation is critical in the maintenance of hepatic differentiation and metabolic zonation by maintaining HNF4α levels.

Additionally, HNF4α is a key event in the development of HCC (7). HCC often manifests as a degenerative phenotype, becomes more severe, and eventually leads to liver failure (7). The pathogenesis of HCC is complex and varies among individuals. One mechanism that we propose that contributes to the development of HCC is loss of O-GlcNAcylation, which leads to loss of HNF4α. Our data are consistent with previous studies in which people with cirrhosis had decreased O-GlcNAcylation compared to healthy individuals (5). This is further sustained in those who develop HCC. However, other studies have shown that O-GlcNAcylation levels can be diverse in people with HCC, suggesting that both increased and decreased levels of O-GlcNAcylation may contribute to the development of HCC in individuals with liver disease (39).

Previous studies on the role of O-GlcNAcylation have used either *in vitro* (3, 4, 39) or xenograft models (3, 39). Our data obtained using cell specific OGT and OGA knockout *in vivo* illustrate that the lack of hepatic O-GlcNAcylation was more severe than increased O-GlcNAcylation. Decreased O-GlcNAcylation led to significant induction in the progression of HCC. Our data show that OGT-KO mice have significant injury, inflammation, fibrosis, disruption of metabolic zonation, and dedifferentiated hepatocytes, all of which are exhibited in HCC progression (38, 40, 41). This could be explained by a multitude of factors. Models knocking out OGT in hepatocytes and biliary cells exhibit an induction of necroptosis, causing liver injury and inflammation (5).

Additionally, O-GlcNAcylation is found to regulate serum response factor (SRF), which leads to the activation of hepatic stellate cells and fibrosis (42). Additionally, our data corroborate other studies showing that O-GlcNAcylation is a critical regulator of cell proliferation (2, 43-45). Further, our data exhibited increased stemness gene expression, indicating a more stem cell-like identity. HNF4α, hepatic master regulator, activity was also significantly downregulated. Both are signs of cell proliferation potential. Multiple pathways in the liver can govern cell proliferation. We found increased activity of YAP signaling, indicating the primary driver of proliferation, with no inductions in β-catenin, ERK, and AKT signaling. The YAP regulation of O-GlcNAcylation is controversial. Two O-GlcNAcylation sites have currently been mapped to YAP, Ser127 and Thr241. Interestingly, one site is thought to inhibit HCC progress (Ser127), while the other enhances disease progression (Thr241) (3, 46). Both modifications act by increasing the translocation and activity of YAP; however, each seems to have a different role. Ser127 leads to increased cell proliferation and survival, whereas Thr241 modification allows YAP to upregulate the transferrin receptor (TFRC), promoting cell death through ferroptosis. One possible explanation for why it contributes to cell proliferation in our models is that HNF4α and YAP activities are intertwined (47-49). One mechanism proposed is that HNF4α competes with YAP in heterodimerization with TEAD4, inhibiting YAP activity (49). This indicates that a lack of HNF4α, triggered due to lack of O-GlcNAcylation, would cause an induction of YAP activity, leading to HCC progression.

In summary, these data show that the loss of O-GlcNAcylation is critical in maintaining hepatic differentiation and liver zonation. Loss of hepatic differentiation and increased cell death further promotes inflammation and fibrosis, ultimately promoting HCC progression. While, increasing hepatic O-GlcNAcylation had no effect on hepatic differentiation or HCC promotion. These data indicate that increasing O-GlcNAcylation could be a novel therapeutic strategy for chronic liver diseases, especially HCC.

## Abbreviations

DC: Dendritic Cells
DEN: diethylnitrosamine
DF: Dedifferentiated
Endo: Endothelial
GlcNAc: N-acetylglucosamine
H&E: hematoxylin and eosin
HCC: Hepatocellular Carcinoma
Heps: Hepatocytes
HNF4α: Hepatocyte Nuclear Factor 4 alpha
HSCs: Hepatic Stellate Cells Institutional Animal Care and Use
IACUC: Committee
IHC: immunohistochemistry
KO: knockout
KUMC: University of Kansas Medical Center
Kup: Kupffer
Monos: Monocytes
MZ: Mid Zonal
NASH: non-alcoholic steatohepatitis
Neut: Neutrophils
NK: Natural Killer
NPCs: nonparenchymal cells
OGA: O-GlcNAcase
OGT: O-GlcNAc Transferase
PAS: Periodic acid–Schiff
PP: Periportal
PTM: post-translational modification
PV: Perivenous
RBC: Red Blood Cell
SEM: standard error of the mean
Ser: Serine
TCGA: The Cancer Genome Atlas
Thr: Threonine
TMA: Tissue Microarray
WT: Wild Type
αSMA: alpha smooth muscle actin

## Acknowledgements

The authors would like to acknowledge BioRender in the aid of developing the graphical abstract. Additionally, the authors would like to thank Huina Cai from the KUMC Pharmacology, Toxicology and Therapeutics Histology Core Laboratory (for her expert assistance throughout these studies) and Clark Bloomer and Rosanne Skinner from the KUMC genomics core (for their assistance with the sequencing).

## Supplementary Figure Legends

**Figure S1.**
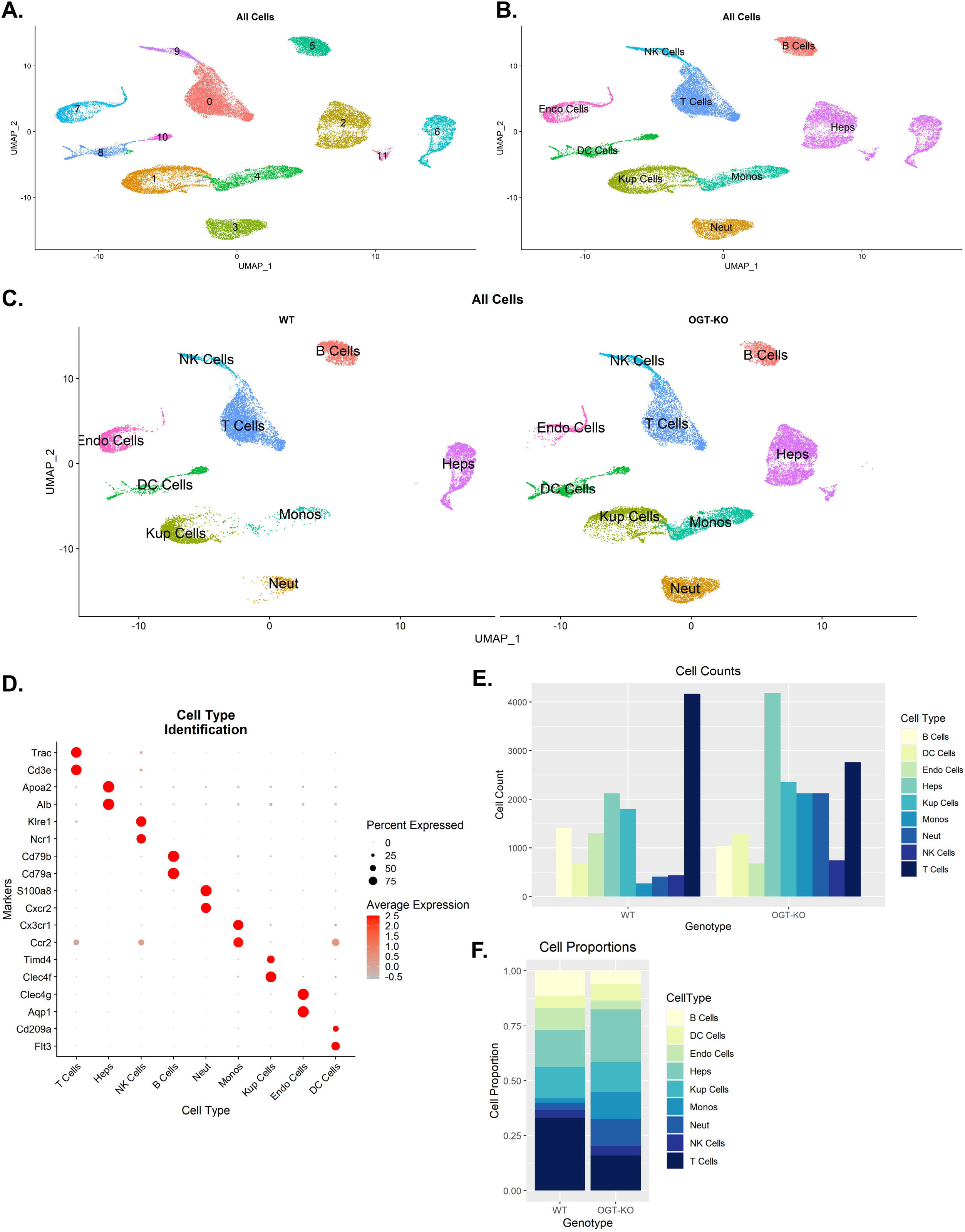
Single-cell RNA-sequencing cell type identification. UMAP of unsupervised clustering with (A) cluster identification number and (B) annotated clusters for OGT-KO and control samples. (C) A split UMAP of annotated clusters between control and OGT-KO. (D) Dot plot showing expression levels and percentage of population expressing two representative markers, which was used to annotate each cell type. Bar graph of (E) total number of cells in each population with (F) respective proportions.

**Figure S2.**
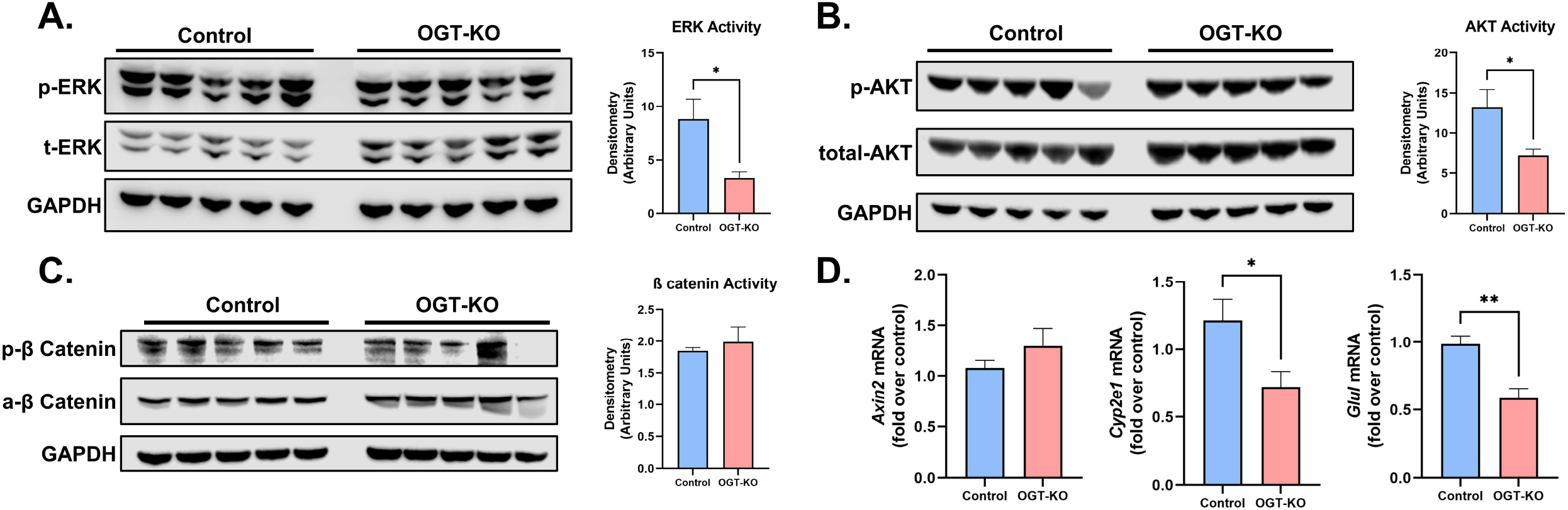
ERK, AKT, p38 and ß-catenin did not contribute to cell proliferation after DEN-induced HCC in OGT-KO mice. Western blot of (A) phosphorylated ERK and total ERK, (B) phosphorylated AKT and total AKT, and (C) phosphorylated ß-catenin and active ß-catenin (non-phosphorylated) with their respective quantification of activity. (D) qPCR of ß-catenin target genes normalized to 18s. Bars represent mean with error bars SEM. Level of significance: **p < 0.01; *p < 0.05 (Two-tailed t-test)

**Figure S3.**
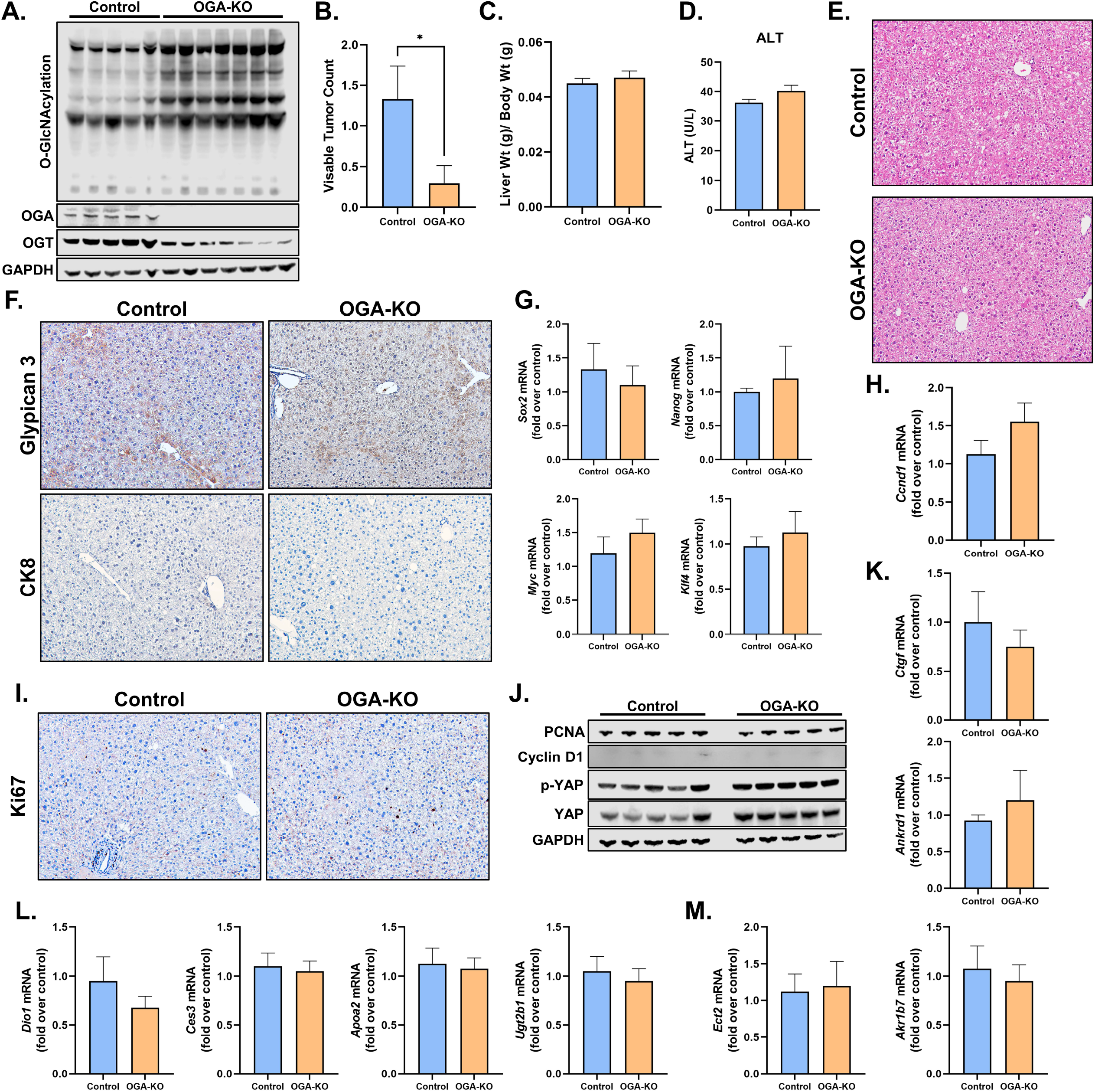
OGA-KO mice showed no significant changes in dedifferentiation and cell proliferation after DEN-induced HCC. (A) Western blot analysis of total hepatic O-GlcNAcylation, OGT, and OGT from OGA-KO and controls treated with DEN. Bar graphs for OGA-KO and control of (B) visible tumor counts, (C) liver-weight-to-body-weight ratio, and (D) serum ALT levels. Photomicrographs of (E) H&E and (F) IHC of HCC markers Glypican 3 and CK8. qPCR of (G) stemness markers (*Sox2, Nanog, Myc*, and *Klf4*) and (H) *Ccnd1*. (I) IHC for the cell proliferation marker Ki67. (J) Western blot analysis of cell proliferation markers PCNA, cyclin D1, YAP, and phosphorylated yap. qPCR of (G) YAP target genes (*Ctgf* and *Ankrd1*) and gene (L) positively (*Dio1, Ces3, Apoa2*, and *Ugt2b1*) and (M) negatively (*Ect2* and *Akr1b7*) regulated by HNF4α. In bar graphs, the bar represents the mean and error bars SEM. Level of significance: *p < 0.05 (Two-tailed t-test)

**Figure S4.**
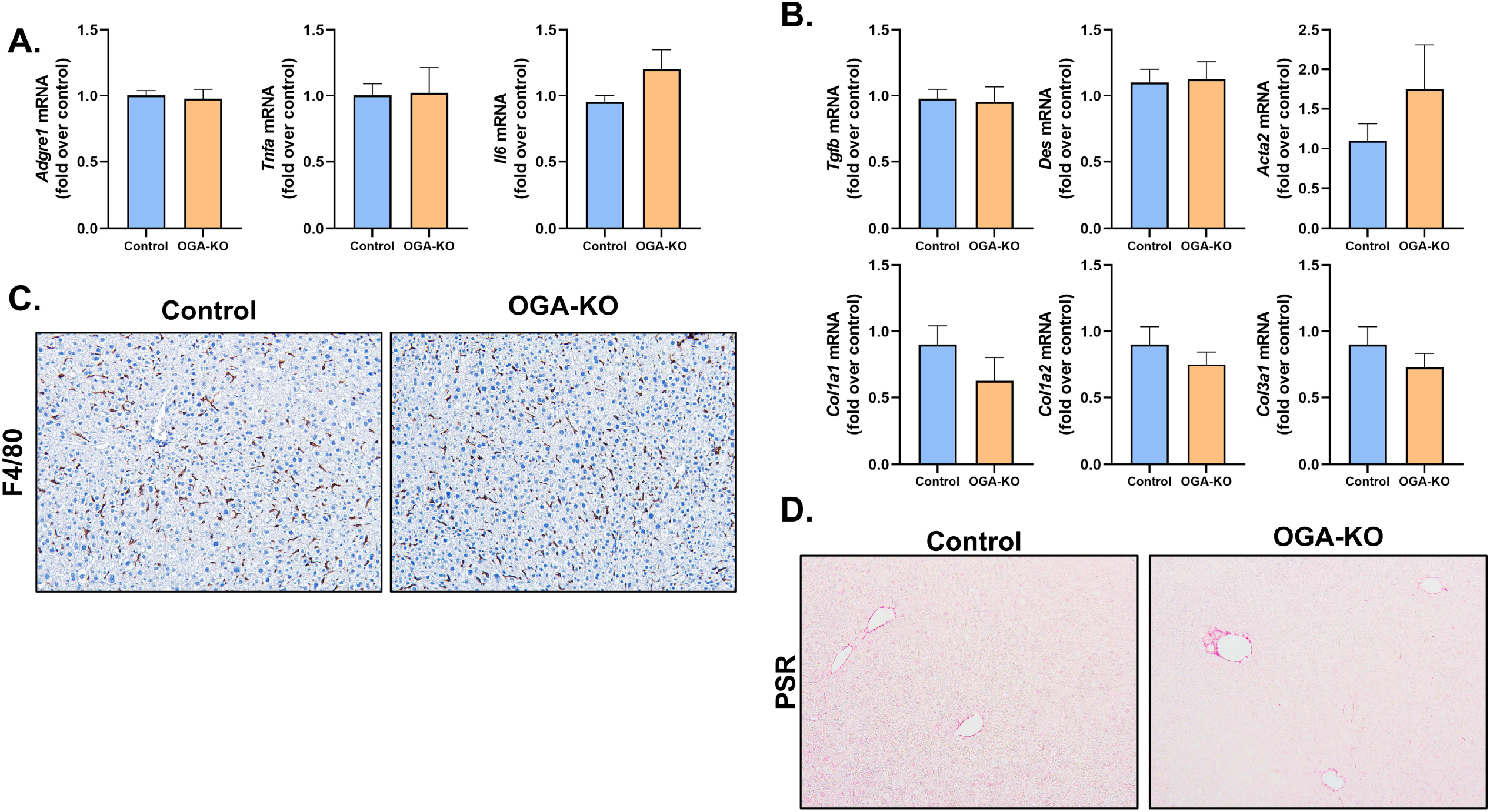
Increased O-GlcNAcylation led to no significant changes in inflammation or fibrosis that were exhibited after DEN-induced HCC. qPCR of (A) proinflammatory markers (*Adgre1, Tnfa*, and *Il6*) and (B) profibrotic genes (*Tgfb1, Des, Acta2, Col1a1, Col1a2*, and *Col3a1*). Bars represent the mean, with error bars meaning SEM. (C) IHC of the macrophage marker F4/80. (D) Picrosirius red staining in the OGA-KO and control mice. Level of significance: (Two-tailed t-test)

